# Prioritizing putatively etiological T cell epitopes across autoimmune diseases

**DOI:** 10.1101/580126

**Authors:** Masato Ogishi

## Abstract

Autoimmune diseases remain a leading cause of mortality among adolescents and young adults worldwide. Despite their clinical impact, there are still significant knowledge gaps in our understanding of immunological tolerance and its breach that characterizes the onset of autoimmune diseases. Genetic associations between the histocompatibility leukocyte antigen (HLA) loci and various autoimmune diseases have been well established. The HLA class I and class II molecules present epitopes to T cells, and T cells play indispensable roles both in the maintenance of tolerance and the pathogenesis of autoimmune diseases. Although a vast number of epitopes and reactive T cell clones have been identified from animal model studies and observational studies, however, only a few have been proven to be causally relevant to disease pathogenesis. Here, we propose a computational framework to prioritize etiologically relevant epitopes by integrating the putatively causal associations between HLA alleles and disease risk identified from population genetics; we define a metric, termed “differential presentation index (DPI),” which principally reflects the relative difference of epitope abundance presented onto HLA molecules whose alleles are genetically predisposing to or protective against the specific disease. We systematically examined publicly available epitope sequence data previously studied in the context of autoimmune diseases. Self-epitopes were generally more stably presented on disease-protective HLAs than non-self epitopes, and hence had a negative DPI. Conversely, proteome-wide sequence alignment revealed that epitopes with highly positive DPI were less similar to self. As a case study, we performed a focused analysis of multiple sclerosis (MS), and identified epitopes from myelin basic protein (MBP), a well-established MS autoantigen, based on DPI-guided prioritization. Moreover, we found several non-MBP-derived self-epitopes with high DPI that are potentially involved in the pathogenesis of MS. Our framework facilitates the identification of etiologically relevant epitopes across autoimmune diseases with known HLA allele association, which in turn expedites the development of epitope-specific disease monitoring and intervention strategies.

## 1 Introduction

Autoimmune diseases have been shown to be more common than previously thought from several epidemiological studies. The estimated total incidence and prevalence are 90 per person-years and 3.2%, respectively, when summed across diseases (1, 2). Graves’ disease (GD), thyroiditis, and rheumatoid arthritis (RA) are among the most common diseases (>10 per 100,000 person-years) whereas other diseases such as systemic lupus erythematosus (SLE) and multiple sclerosis (MS) are relatively rare (1∼10 per 100,000 person-years). Although clinical manifestations considerably vary between diseases, most of the tissue/organ damage is thought to arise from the dysregulated response of the adaptive immunity, both cytotoxic and humoral immunity, to self-antigens. Although significant advances have been made both in the diagnosis and clinical management in the past decade, in most of the diseases there remains no successful therapeutics leading to complete remission and cure. Patients tend to suffer from the chronic, relapsing and refractory nature of the diseases and the adverse effects of treatments, leading to poor quality of life and significant economic burden. Indeed, autoimmune diseases are still among the leading causes of mortality among young and middle-aged adults, particularly among females, and the mortality rate remains relatively constant (3, 4).

Very little is known about the disease-initiating processes of autoimmune diseases despite the advances in our understanding of their steady-state pathophysiology. Mechanistic studies have been hampered, at least in part, because of the following reasons: (i) patients with an early stage disease lacking typical clinical manifestations tend not to seek a medical examination or to receive a correct diagnosis, and therefore are rarely studied; (ii) it is often difficult to elucidate the mechanisms essential for disease prevention and/or initiation from observational studies of patients with chronic inflammation and extensive systemic involvements due to several secondary changes; (iii) etiologies identified from studies utilizing genetically predisposing animal models do not necessarily reflect the pathophysiology in humans. Undoubtedly, identification of epitopes relevant to the disease initiation process would have significant translational implications, since recent studies have indicated the possibility of antigen/epitope-specific immunosuppression via various strategies such as transplantation of receptor-engineered T cells with immunomodulatory functions, and more interestingly, systemic administration of peptide-MHC-based nanomedicine to reconstruct physiological regulatory networks (5–9).

Genetics, likewise external environmental factors, is apparently one of the most upstream factors in the cascade of disease initiation. In other words, the genome alteration can be causal to disease, but not *vice versa* (except tumorigenesis). Notably, a number of major histocompatibility complex [MHC; also known human leukocyte antigen (HLA) in humans] loci, as well as hundreds of non-MHC loci, have been associated with predisposition and protection of autoimmune diseases (10– 12). Since experimental genetic manipulation is not ethically acceptable, population genetics is a valuable tool to assess the contribution of particular HLA alleles to the onset of autoimmune diseases in humans. In other words, the enrichment or paucity of specific HLA alleles in patients with specific autoimmune diseases could reflect their etiological roles, although one should keep in mind the caveat that other non-HLA loci in strong linkage disequilibrium are of *bona fide* etiological significance. That being said, considering the biological function of MHC molecules to present a short peptide fragment (called epitope) derived from various self-and non-self-antigens to T cells, and the multifaceted roles of T cells as the master regulator of adaptive immunity and self-tolerance, the contribution of T cell epitope to autoimmune diseases has been extensively studied, and numerous self-and non-self-derived epitopes and cognate cross-reactive T cell clones have been identified (13–15). Nevertheless, only a minority of them have been causally linked to either disease initiation by pathogenic T cells or maintenance of self-tolerance by clonal deletion at thymus and/or induction of regulatory T cells (Tregs) *in vivo*, and consequently, there remain virtually no epitope-specific diagnostic/therapeutic strategies clinically available to date. Therefore, an additional “filter” complementary to experimental validation of T cell recognition *in vitro* is needed to expedite the exploration of the most etiologically responsible epitopes.

In this context, here we propose a simple approach to prioritize epitopes with high pathophysiological relevance by integrating population genetics and MHC binding prediction. We hypothesized that epitopes more stably presented on the HLA molecules encoded by disease-predisposing alleles are more likely to be pathogenic, whereas those preferentially presented on HLA molecules encoded by disease-protective alleles are more likely to contribute to tolerance. HLA loci associated with specific diseases were extracted from the previous phenome-wide association study (PheWAS) data (11), and MHC binding prediction was conducted bioinformatically using NetMHCpan and NetMHCIIpan (16, 17). We introduced a metric termed “differential presentation index (DPI)” based on the predicted binding strength among predisposing and protective HLA molecules. As a case study, we screened epitopes studied in the context of MS, and, as expected, found that several epitopes derived from myelin basic protein (MBP), a well-characterized autoantigen, ranked highly based on MS-specific DPI (18, 19). Moreover, we found candidates of MS-relevant non-MBP epitopes derived from self-antigens including SIK1, GRK2, IFNB, and EPO, all of which are present and play critical roles in the central nervous system (CNS). Notably, some of those newly identified epitopes had even higher MS-specific DPI than MBP epitopes. Finally, examination of an experimentally verified molecular mimicry epitope dataset revealed two putatively MS-predisposing mycobacterial epitopes homologous to an MBP-derived self-epitope and one putatively RA-protective mycobacterial epitope homologous to mammalian 60kDa heat shock protein (HSP60), a well-known Treg-inducing self-antigen (20, 21). Collectively, these findings illustrate the utility of DPI-based epitope prioritization strategy in search of etiologically relevant epitopes. Further characterization in different disease contexts and experimental validation are warranted. The datasets and codes necessary to reproduce the analytical pipeline are made publicly available as the R package *DPA* on GitHub (https://github.com/masato-ogishi/DPA/) to expedite future research.

## 2 Results

### 2.1 Datasets

We extracted disease-HLA associations across different autoimmune diseases from the previous study (11). This study involved two populations of European ancestry individuals (N=28,839 and 8,431) and tested the association of HLA variation with 1,368 phenotypes. We identified the following autoimmune disease phenotypes: ankylosing spondylitis (AS), celiac disease (CD), dermatomyositis (DM), giant cell arteritis (GCA), Graves’ disease (GD), Juvenile rheumatoid arthritis (JRA), localized lupus and systemic lupus erythematosus (SLE), multiple sclerosis (MS), polymyalgia rheumatica (PMR), polymyositis (PM), primary biliary cirrhosis (PBC), psoriasis and psoriatic arthropathy (PSO), rheumatoid arthritis (RA), systemic sclerosis (SS), type 1 diabetes (T1D), ulcerative colitis (UC), and Wegener’s granulomatosis (GPA). We classified the associated HLA alleles into two categories, namely, “predisposing” and “protective,” based on the odds ratios. Then we screened diseases that have both predisposing and protective HLA alleles. The following diseases met the criteria: AS, CD, DM, GD, JRA, SLE, MS, PMR, PBC, PSO, RA, SS, T1D, and UC (Table S1). Next, we downloaded the linear T cell epitope data annotated with at least one cell-based functional assay results from the Immune Epitope Database (IEDB) (22). Inclusion/exclusion criteria in terms of assay annotations are provided in Table S2. Epitopes were considered “immunogenic” if a positive T cell response was recorded from at least one functional assay. We use this definition to capture as many potentially T-cell-activating epitopes. We identified epitopes studied in the context of the following autoimmune diseases: AS, CD, GD, GPA, MS, PBC, PSO, RA, SLE, SS, and T1D (Table S3). It should be noted that these disease contexts do not necessarily guarantee the pathophysiological relevance of the epitopes studied. We screened exact matches to human proteome (UniProt ID: UP000005640) to identify self-derived (S) and non-self (NS) epitopes. We also aligned each epitope sequence against the entire human proteome using the Smith-Waterman local alignment algorithm, with a substitution matrix and gap opening/extension penalty parameters identical to those utilized in the blastp-short tool (see Materials and Methods). We took three representative metrics, namely, mean, maximum, and minimum, of the alignment score distribution as indicators of similarity-to-self for each of the epitopes.

### 2.2 Differential presentation analysis

Our goal was to identify epitopes the most differentially presented among predisposing and protective HLA molecules in a disease-specific context (Figure 1). To this end, we first merged the disease-HLA and disease-epitope association data to obtain a set of epitopes coupled with disease-specific predisposing/protective HLA allele information. Then, we computed the percentile rank values using either NetMHCpan or NetMHCIIpan. Only four-digit HLA alleles were considered, and two-digit HLA alleles were ignored in the subsequent analysis. Computation for HLA-DP/DQ alleles was problematic because a DP-DQ pair must be provided for the binding prediction, despite the single allelic format of disease association in the PheWAS source data. We ended up computing every single possible combination of DP-DQ alleles that contain at least one disease-associated allele and summarizing the predicted values by taking their medians. We then summarized the sign-inverted log-transformed percentile rank values among protective and predisposing HLAs by taking their mean, maximum and minimum. Note that, after this conversion, the larger value reflects more stable binding. We defined “differential presentation index (DPI)” as the difference between the maximum values among predisposing and protective HLAs. We utilized the maximum value among HLAs tested because any epitope does not necessarily bind strongly to all disease-associated HLAs; strong binding to at least one disease-associated HLA is adequate as the initial inclusion criteria. DPI is an indicator unique to each epitope defined in a disease-specific context. A positive DPI means that the epitope is predicted to be bound more strongly to at least one of the predisposing HLAs than any of the protective HLAs. We then categorized epitopes with DPI > 0.5 and DPI < −0.5 as “predisposing” and “protective,” respectively. Note that these epitope-level categories are putative. We excluded epitopes if the minimum percentile rank among all disease-associated HLAs was higher than the recommended threshold (2% and 10% for HLA-I and HLA-II, respectively). Figure 2 shows the distributions of the lowest percentiles among predisposing and protective HLAs. Both the number of available epitope data and their distribution considerably varied between diseases. There was a relatively large number of epitopes associated with MS, RA, and T1D (Figure 2B). Full epitope data is summarized in Table S4.

**Figure 1.**
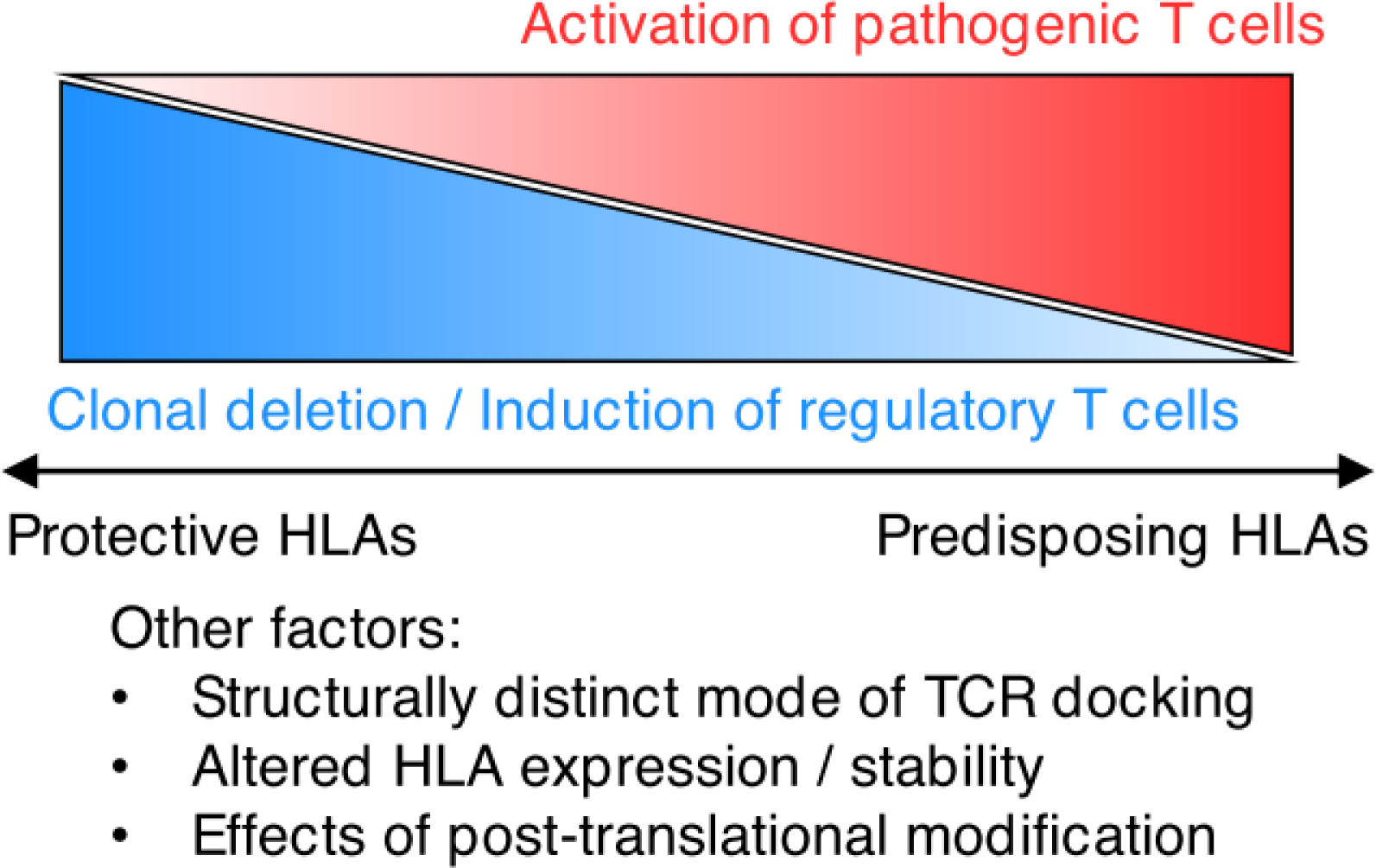
A diagram of differential peptide presentation in association with autoimmunity. Several genetic associations between HLA alleles and various autoimmune diseases have been identified. Given the biological function of HLA molecules, we hypothesized that HLA molecules whose alleles are genetically associated with disease predisposition presumably present either pathogenic epitopes more or protective epitopes less, and likewise, HLA molecules whose alleles are genetically associated with disease protection presumably present either pathogenic epitopes less or protective epitopes more. Note that this scheme is oversimplified from the following viewpoints; first, the genetic association with HLA loci could result from other etiologically responsible loci in linkage disequilibrium; second, the different contribution to disease pathogenesis between HLA alleles could be explained in multiple ways other than differential epitope presentation (12); lastly, although in this study we use the percentile rank values predicted by either NetMHCpan or NetMHCIIpan as a surrogate of the stability of epitope presentation, neither affinity or affinity-based percentile rank is the only parameter representing the HLA-epitope thermodynamic interaction.

**Figure 2.**
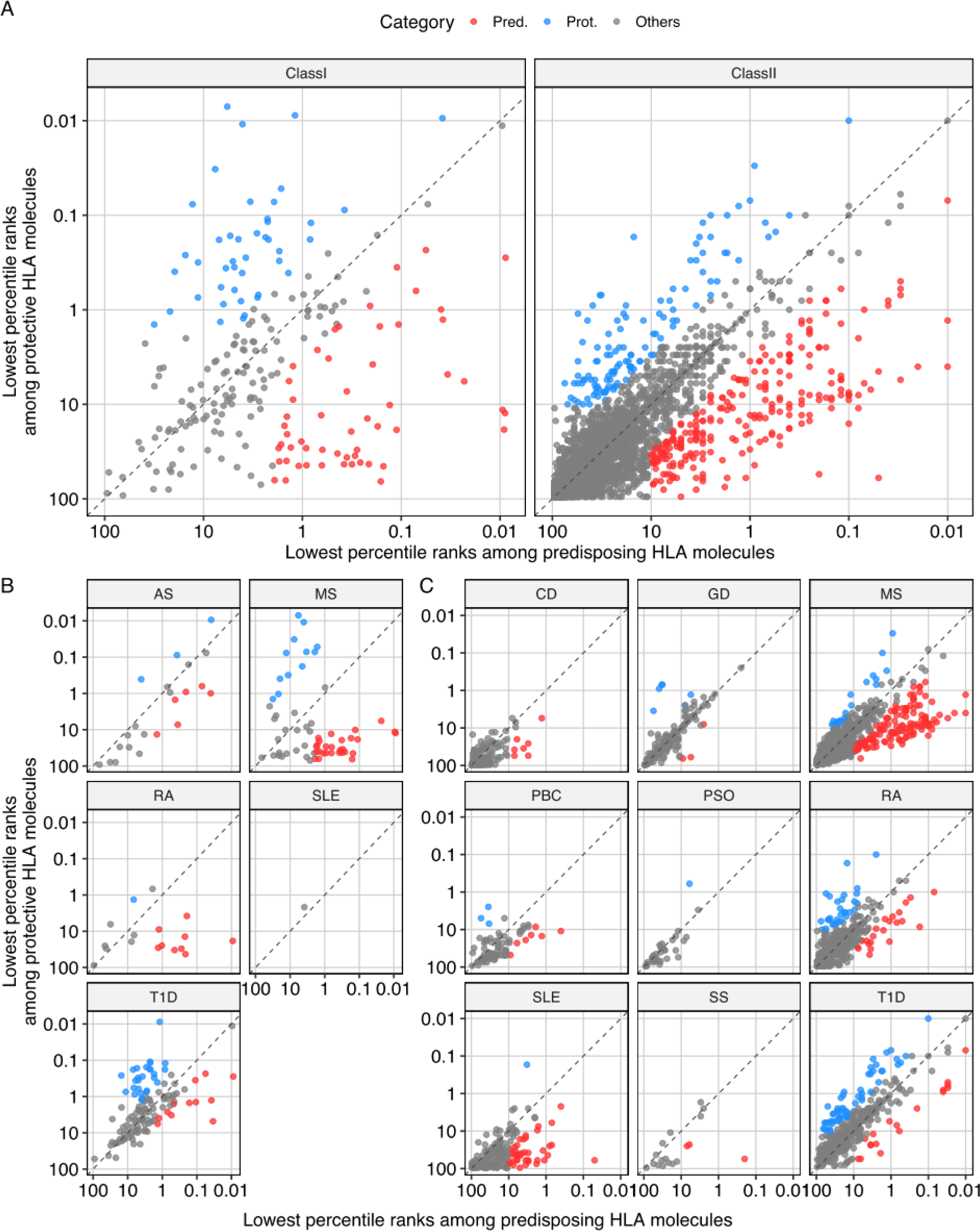
Differentially presented T cell epitopes studied in the context of autoimmune diseases. (A) Differential presentation analysis of autoimmunity-associated HLA-I and HLA-II epitopes. The axes represent the lowest percentile rank values (*i.e.*, the most stable binding) among disease-predisposing and disease-protective HLA alleles. Epitopes were categorized into putatively disease-predisposing (Pred.), putatively disease-protective (Prot.), or others based on the following criteria: (i) the lowest percentile rank among HLA alleles tested was below the HLA class-specific threshold (*i.e.*, 2% and 10% in HLA-I and HLA-II epitopes, respectively); DPI was either higher than 0.5 or lower than −0.5. The DPI threshold of 0.5 was arbitrarily determined, which roughly corresponds to a three-fold change in the percentile rank. We utilized the lowest percentile rank among disease-associated HLA alleles as a representative metric to capture any epitope bound stably to at least one of the disease-associated HLAs as a potentially etiologically relevant epitope. (B and C) Differential presentation analysis of (B) HLA-I and (C) HLA-II epitopes, stratified by the autoimmune diseases in which the epitopes have been studied. AS, ankylosing spondylitis. CD, celiac disease. GD, Graves’ disease. MS, multiple sclerosis. PBC, primary biliary cirrhosis. PSO, psoriasis. RA, rheumatoid arthritis. SLE, systemic lupus erythematosus. T1D, type 1 diabetes.

Next, we explored the common characteristics among the differentially presented epitopes. Interestingly, predisposing epitopes were significantly more likely to be derived from non-self antigens than protective epitopes (*P*=3.2×10^−6^ and 4.7×10^−7^ in HLA-I and HLA-II, respectively, by chi-square test) (Figure 3A). Meanwhile, we noted that 31/55 (56%) of HLA-I predisposing epitopes were classified as non-immunogenic, meaning that there was not even a single positive T cell assay result (Figure 3B). The difference in terms of immunogenicity between predisposing and protective epitopes was also statistically significant in HLA-I but not in HLA-II epitopes (*P*=2.5×10^−7^ and 0.35, respectively, by chi-square test). To quantitatively assess similarity-to-self, we chose the highest sequence alignment scores, representing the best-match sequence-level homology against the human proteome. As expected, epitopes exactly matched to human proteome had apparently higher scores (Figure 3C). Notably, the predisposing epitopes were less similar to self (*P*=1.6×10^−6^ and 1.1×10^−6^ in HLA-I and HLA-II, respectively, by Wilcoxon rank sum test) (Figure 3D). Consistently, the overall correlation between DPI and similarity-to-self was evident, although we noted considerable variation between diseases (Figure S1). Of note, MS-associated predisposing epitopes with high DPI were more non-self, an observation not contradictory to the molecular mimicry hypothesis (13, 23, 24). In contrast, no correlation was observed among T1D-associated epitopes, possibly indicating an indispensable role of aberrantly formed self-epitopes (25, 26). It is difficult to interpret epitopes associated with other diseases due to the paucity of data. Type 2 ANOVA revealed that neither peptide category or T cell reactivity significantly contributed to the similarity-to-self variation after stratified by peptide origin (S vs. NS) in HLA-I, whereas mutual interaction between peptide category and origin existed (*P*=2.7×10^−9^) in HLA-II (Figures 3E and F). It was unexpected to us that T cell recognition was not associated with the similarity-to-self score. We did not either observe any difference among immunogenic epitopes and non-immunogenic MHC binders in the datasets we have previously compiled (N=21,162 and 31,693 for HLA-I and HLA-II epitopes, respectively) (Figure S2) (27) (manuscript under review in *Frontiers in Immunology*).

**Figure 3.**
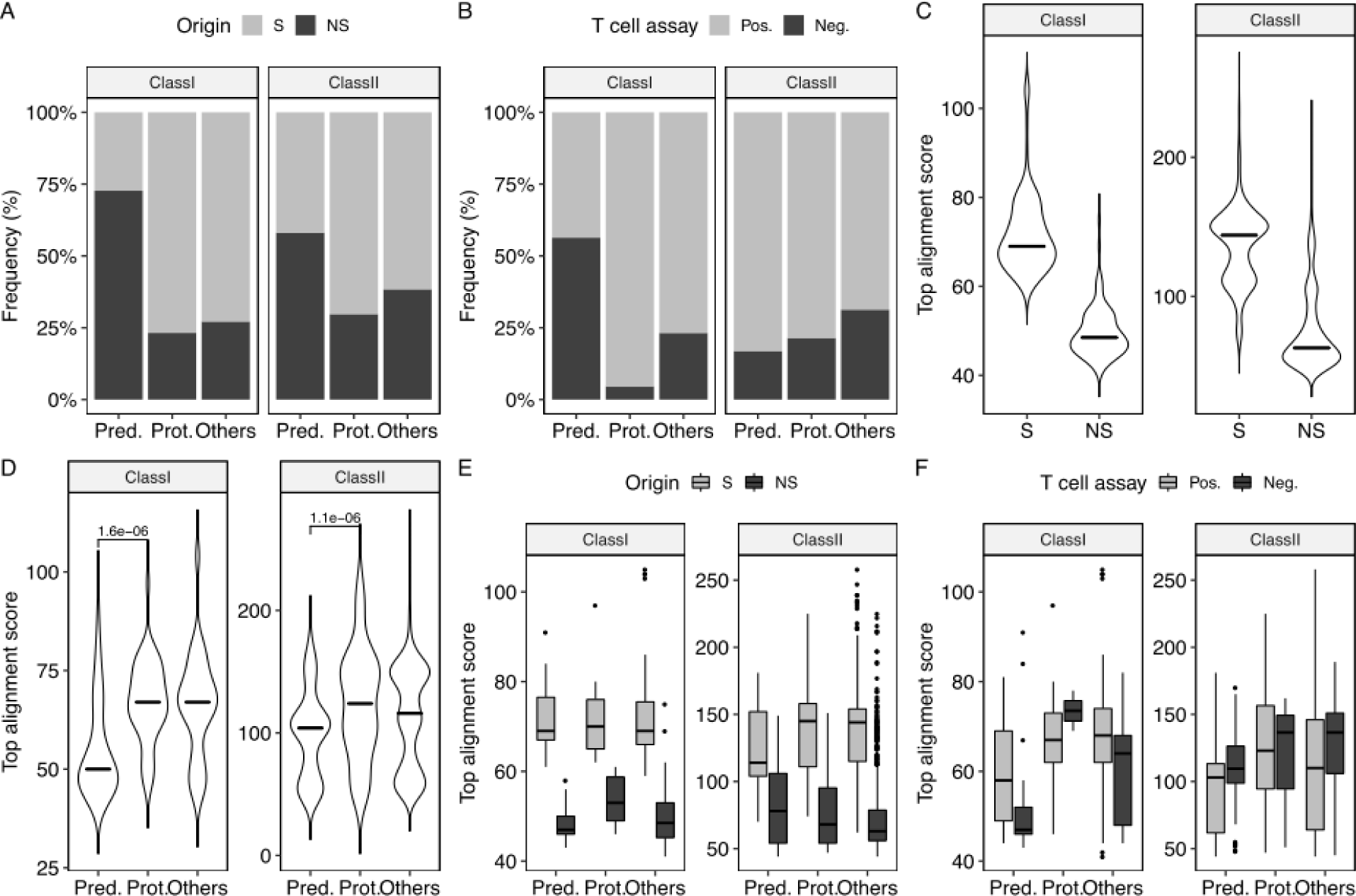
Analysis of differentially presented epitopes across autoimmune diseases. (A) Associations between the differential presentation categories and origins of the epitopes (N=220 and 2520 for HLA-I and HLA-II, respectively). Epitopes with and without at least one exact sequence match in the human proteome (UniProt ID: UP000005640) were considered self (S) and non-self (NS), respectively. (B) Associations between the differential presentation categories and the epitope immunogenicity determined by IEDB-derived annotations of functional T cell assay results. Epitopes with at least one positive assay result were considered positive (Pos.), and those only having negative assay results were considered negative (Neg.). (C-F) Distributions of the top sequence alignment scores against the entire human proteome as a surrogate indicator of similarity to self. Local sequence alignment was conducted by employing the Smith-Waterman algorithm with the substitution matrix and gap-opening/extension costs identical to those used in the blastp-short program. Statistical significance was determined by Wilcoxon’s signed rank test.

In summary, (i) predisposing epitopes are generally less similar to self, primarily because peptides derived from non-human antigens tend to be more stably presented on predisposing HLAs; (ii) predisposing HLA-I but not HLA-II epitopes are less likely to be recognized by T cells; and (iii) propensity of T cell recognition is orthogonal to peptide similarity-to-self.

### 2.3 Case study: multiple sclerosis

We next asked whether the proposed DPI-based framework was able to effectively prioritize known antigens/epitopes. To examine this, we decided to do a focused analysis of MS-associated epitopes because of the abundance of available epitope data. DPIs were calculated based on the binding prediction to four and six HLA-I and HLA-II alleles associated with MS, among which two and three were MS-predisposing, respectively. More than 75% of both HLA-I and HLA-II MS-predisposing epitopes did not match to the human proteome (*P*=2.3×10^−4^ and 1.1×10^−5^ in HLA-I and HLA-II, respectively, by Fisher’s exact test) (Figure 4A). Meanwhile, there was a striking dissociation in terms of T cell recognition between HLA-I and HLA-II MS-predisposing epitopes (*P*=6.0×10^−8^ and 1.0×10^−5^ in HLA-I and HLA-II, respectively, by Fisher’s exact test) (Figure 4B). Interestingly, most of the non-self, non-immunogenic, yet predisposing HLA-I epitopes were derived from human endogenous retrovirus W (HERV-W). Predisposing epitopes had lower similarity-to-self compared to protective epitopes in HLA-I but not in HLA-II data (*P*=4×10^−5^ and 0.07 in HLA-I and HLA-II, respectively, by Wilcoxon rank sum test) (Figure 4C). We speculated that this is owing to a set of predisposing HLA-II epitopes highly homologous to MBP-derived epitopes. As expected, a subset of non-MBP-derived predisposing HLA-II epitopes shared high sequence homology to MBP (*P*=0.3 and 1×10^−4^ in HLA-I and HLA-II, respectively, by Wilcoxon rank sum test) (Figure 4D). Finally, we found that DPI-guided prioritization enriched MBP-derived epitopes among self-epitopes (*P*=0.01 and 1×10^−4^ in HLA-I and HLA-II, respectively, by Wilcoxon rank sum test) (Figure 4E).

**Figure 4.**
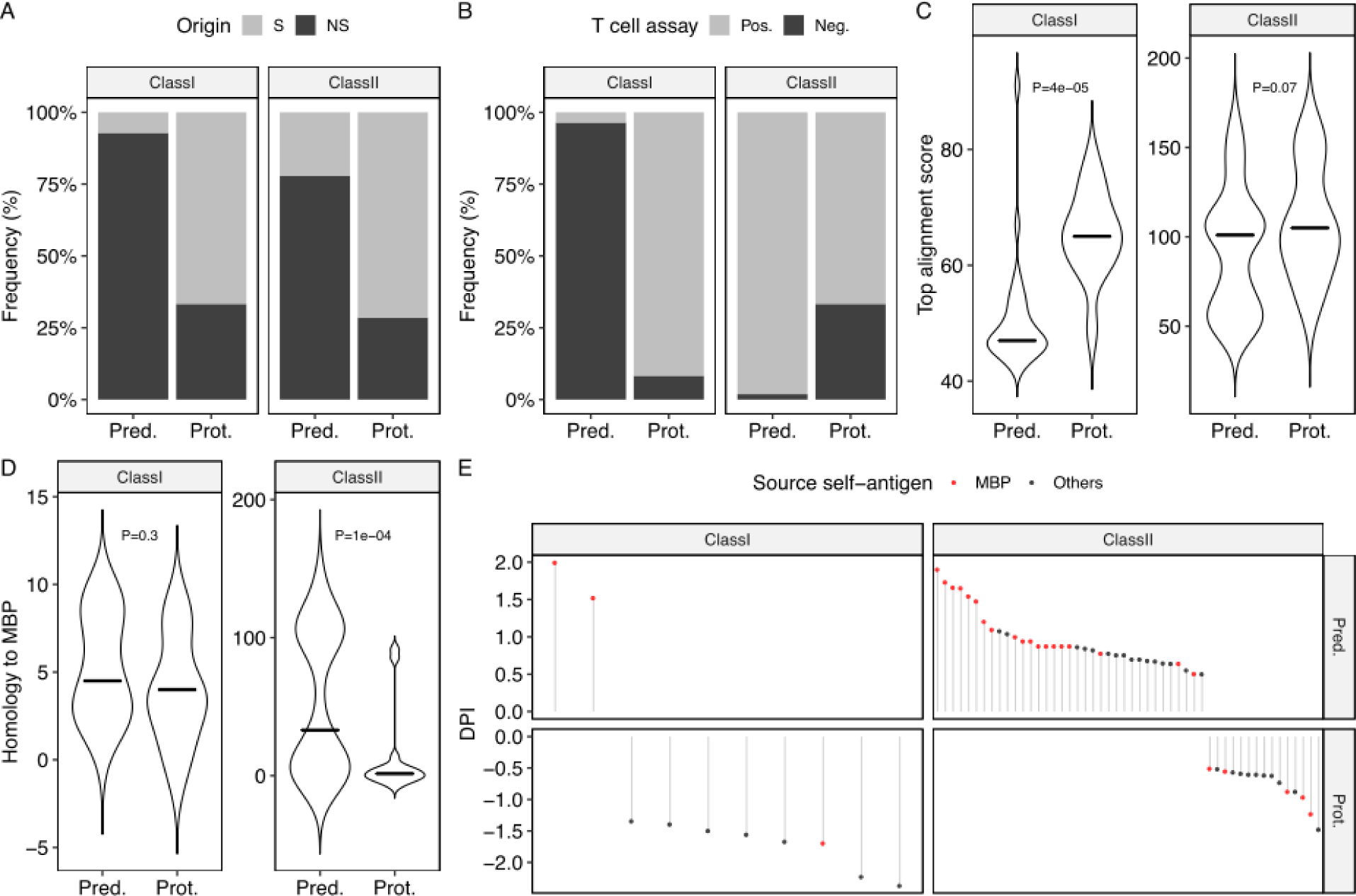
Analysis of differentially presented epitopes associated with multiple sclerosis. (A-B) (A) Origins and (B) T cell assay results of the MS-associated epitopes (N=62 and 907 for HLA-I and HLA-II, respectively). (C and D) Distributions of the top sequence alignment scores against (C) the entire human proteome and (D) a human MBP protein (UniProt ID: P02686). Statistical significance was determined by Wilcoxon’s signed rank test. (E) Prioritization of epitopes with putative etiological relevance based on their DPIs. Epitopes with exact sequence matches to human MBP are shown in red.

Encouraged by these observations, we next sought to screen epitopes potentially relevant to MS etiology. We applied our DPI calculation framework to all self-epitopes identified from the previously compiled epitope datasets (Table S5) (27). The top five putatively MS-predisposing epitopes for both HLA-I and HLA-II are shown in Table 1. Interestingly, the top-ranked HLA-I epitope RPRPVSPSSL is derived from salt-inducible kinase1 (SIK1). The fourth HLA-I epitope KPRSPVVEL is derived from G-protein coupled receptor kinases 2 (GRK2). Notably, the top-ranked HLA-I epitopes had even higher MS-specific DPI than MBP-derived epitopes. Likewise, among HLA-II epitopes, two interferon beta (IFN-β)-and two erythropoietin (EPO)-derived epitopes were identified (Table 1). MBP-derived HLA-II epitopes were excluded from Table 1 but can be found in Table S5. IFN-β has traditionally been used as an immunomodulatory drug for MS, and is thought to act as a suppressor for T helper type 1 (Th1) cells and T helper 17 (Th17) cells (28–30). EPO, although initially identified as an essential factor for hematopoiesis secreted from kidney, has been implicated as a potent neuroprotective agent (31). In addition to MS-predisposing epitopes, we also explored the putatively MS-protective epitopes. The top five putatively MS-protective HLA-II epitopes are shown in Table 2. Of note, two glutamic acid decarboxylase (GAD) epitopes were identified. Note that we labeled epitopes as “disease-protective” based on genetic association, and hence it does not necessarily guarantee their role in the maintenance of self-tolerance (*e.g.*, induction of Tregs). Nevertheless, it is possible that at least some epitopes with considerably low DPIs act as Treg-inducing epitopes. We will discuss this point in the next chapter.

**Table 1.**
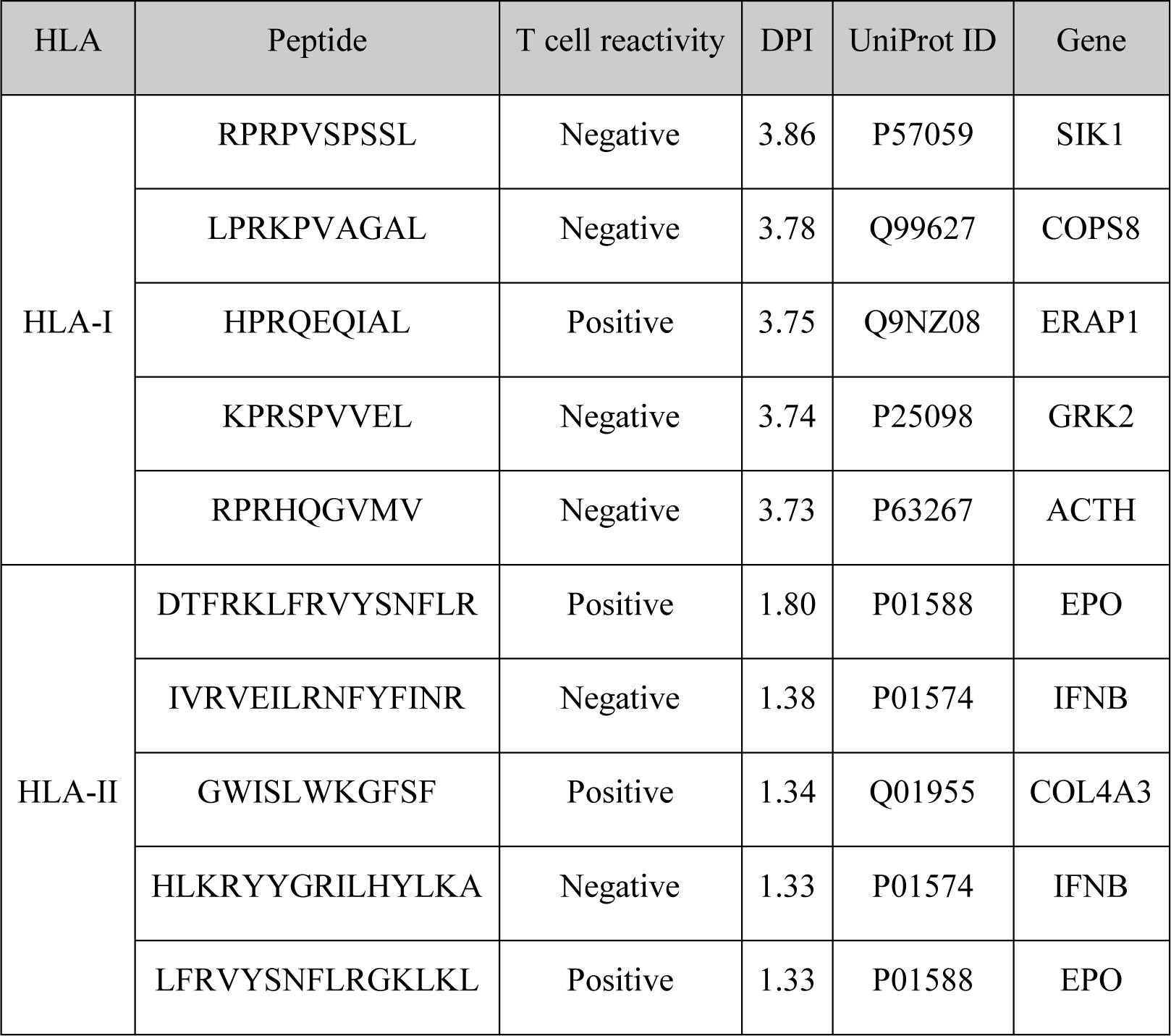
Putatively MS-predisposing non-MBP-derived self-epitopes. DPI was calculated based on the predicted binding to MS-associated HLAs. Top five epitopes were shown. A complete list of epitopes examined can be found in Table S5.

**Table 2.** Putatively MS-protective non-MBP-derived self-epitopes. DPI was calculated based on the predicted binding to MS-associated HLAs. Top five epitopes were shown. A complete list of epitopes examined can be found in Table S5.

| HLA | Peptide | T cell reactivity | DPI | UniProt ID | Gene |
| --- | --- | --- | --- | --- | --- |
| HLA-II | GWYTYMLVPAALTGL | Negative | -1.80 | A1A5B4 | ANO9 |
|  | NFFRMVISNPA | Positive | -1.60 | Q05329 | GAD2 |
|  | QMTFLRLLSASAHQN | Positive | -1.60 | P20908 | COL5A1 |
|  | FFRMVISNPAATHQD | Positive | -1.54 | Q05329 | GAD2 |
|  | FLKKFHFLKGATLC | Positive | -1.48 | Q4ZG55 | GREB1 |

### 2.4 Case study: molecular mimicry

Molecular mimicry has been proposed as one of the potential mechanisms underlying the breach of self-tolerance and autoimmunity (23, 32–34). However, the pathophysiological significance of the T cell clones recognizing both self-and pathogen-derived epitopes *in vivo* remains undetermined or inconclusive in most cases. Moreover, recognition of a specific epitope by effector T cells *in vitro* does not preclude the possibility of recognition of the same epitope by Tregs *in vivo*, which may overall result in the protection against disease.

Self-and pathogen-derived epitopes associated with diseases in the same direction are likely to share biological roles as well as sequence-level homology. Thus, we searched pairs of human-pathogen epitopes that have the differential presentation category in common. We utilized the miPepBase, a database of experimentally verified self-epitopes and mimicking pathogen-derived epitopes (35). Surprisingly, among the forty-three epitope pairs identified, only four had the same differential presentation category in common (Table 3). Of note, there were two mycobacterium epitopes associated with MS. The role of mycobacteria in the pathogenesis of MS remains controversial (36), and therefore the two mycobacterial epitopes and the corresponding self-epitope appear to be worth being further investigated. Meanwhile, there was one epitope pair associated with putative RA protection. Interestingly, both epitopes are derived from heat shock protein (HSP). It is known that cell stress-induced up-regulation of HSPs and the resultant presentation of HSP epitopes induce Tregs (37). Indeed, T cell clones cross-reactive to epitopes derived from mammalian HSP60 and *M. bovis* HSP65 have been shown to protect against experimental arthritis, strongly suggesting the involvement of Tregs (20, 38). The comprehensive set of epitope pairs tested are summarized in Table S6.

**Table 3.**
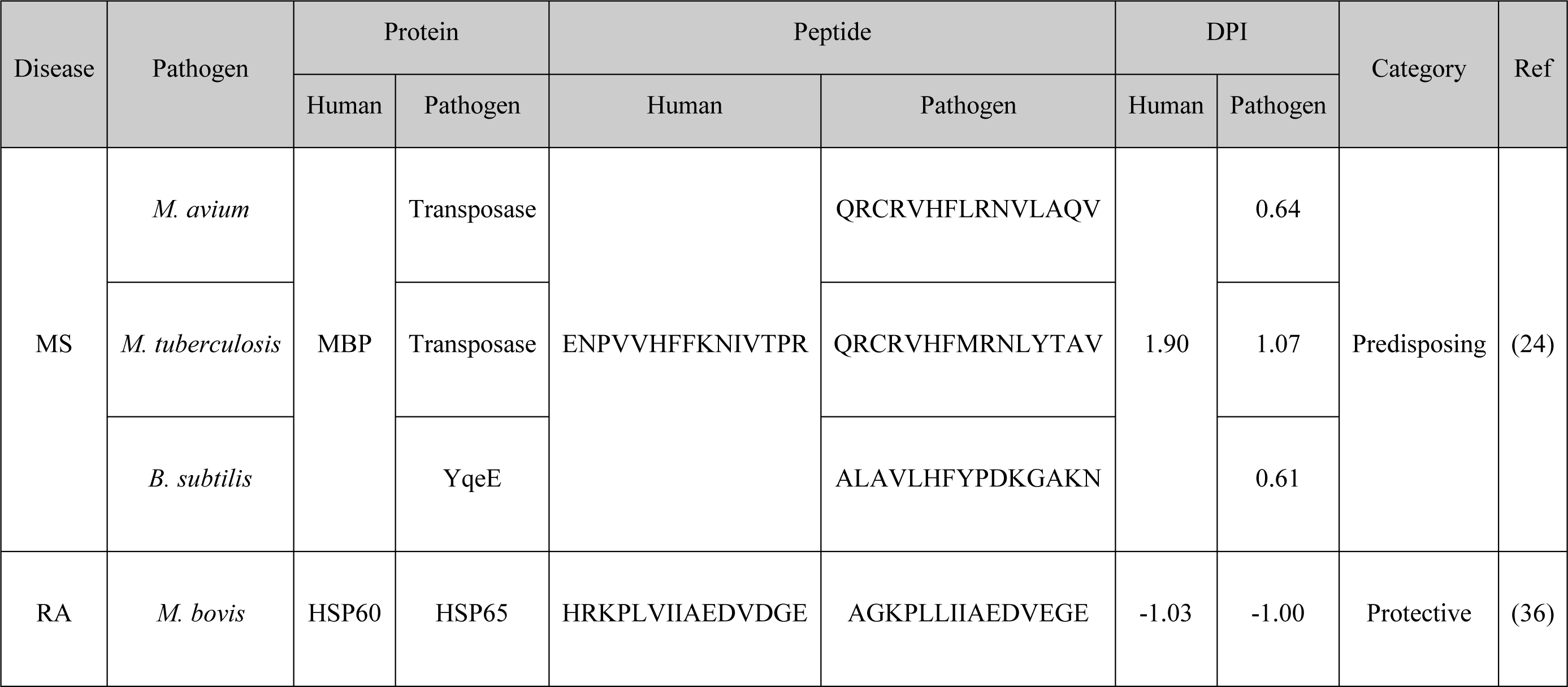
Human-and pathogen-derived HLA-II epitope pairs having the same differential presentation category in common. A complete list of epitopes examined can be found in Table S6.

## 3 Discussion

We systematically mined several T cell epitope candidates likely to be etiologically relevant across autoimmune diseases by differential presentation onto genetically inferred disease-predisposing and -protective HLA molecules. T cells have been thought to play multifaceted roles both in the maintenance of physiological self-tolerance and in autoimmune diseases. To date, T cells epitopes presented on HLA molecules whose alleles are genetically associated with disease predisposition have been extensively studied. However, to our knowledge, there are yet few studies systematically comparing epitope presentation on both predisposing and protective HLAs. Probably, one reason is that testing multiple HLA alleles experimentally is labor-intensive and cost-prohibitive. For example, there are 38 HLA alleles significantly associated with T1D based on the PheWAS data we utilized in this study. Instead of experimental testing, we conducted a comparative analysis through a bioinformatic approach, which can be adequate for a screening purpose. The DPI-based classification system of putatively predisposing and protective epitopes proposed in this study is a novel approach complementary to the current standard approach, *i.e.*, detection of cognate T cell clones. We used percentile rank for differential presentation analysis because this metric is not affected by inherent bias of specific HLA molecules towards higher or lower mean predicted affinities and thus allows a direct comparison between different HLA molecules (16). We used the lowest percentile rank values among predisposing and protective HLA molecules for determining the degree of differential presentation. This definition is conceptually equivalent to the comparison between two hypothetical populations having either all disease-specific predisposing or protective HLA alleles (instead of six HLA alleles per individual in the real-world). We did not consider the odds ratio of different HLA alleles for a specific disease. For example, the odds ratios among T1D-and MS-predisposing HLA-II alleles ranges from 4.420 to 1.713 and 2.672 to 3.475, respectively. Whether odds-ratio-weighted metric contributes to better prioritization of etiologically relevant epitope may be a topic of further investigation.

One of the advantages of computational screening is its unbiasedness; experimental approach often focuses on either known epitopes or epitopes from known antigens, although this type of investigation is likely to overlook epitopes of *bona fide* etiological significance derived from unknown antigens. We tackled this issue by performing an unbiased screen of epitope candidates differentially presented onto MS-predisposing and -protective HLA molecules as a case study. As expected, our analysis revealed several MBP-derived epitopes. The current consensus is that MS is a T cell-mediated autoimmunity against myelinated self-antigens including MBP, and the etiological significance of MBP in MS has been well documented (18, 39). Therefore, this enrichment of MBP-derived epitopes can be viewed as an internal positive control for the analysis. Furthermore, we identified several self-antigens, including SIK1, GRK2, IFNB, and EPO, as a potential source of etiologically relevant epitopes. First, SIK1 mutations have been documented as a cause of severe developmental epilepsy (40, 41). Second, a previous study of experimental autoimmune encephalomyelitis, a mouse model for MS, has shown that GRK2^+/-^ mice expressing 50% of the GRK2 protein did not suffer from relapses unlike the wild-type animals, and the absence of relapse was associated with a marked reduction of infiltrating inflammatory cells in the CNS (42). Modulation of the Toll-like receptor signaling via GRK2 in microglia was also reported (43). Third, IFN-β is a naturally occurring cytokine mediating a wide range of anti-inflammatory responses in the CNS, and has been used as a therapeutic agent for MS (28, 30). The formation of neutralizing autoantibody against therapeutically administered IFN-β strongly suggests a pre-existing humoral immune response against this self-antigen (44). Fourth, EPO is an endogenous neuroprotective protein, and its efficacy has been shown in a pre-clinical model and a small clinical trial, although the subsequent study failed to show its superiority (31, 45, 46). Besides the putatively MS-predisposing self-antigens, we also identified putatively MS-protective candidate self-antigens including GAD. GAD catalyzes the synthesis of gamma-aminobutyric acid (GABA), an inhibitory neurotransmitter, from glutamate, and hence has an indispensable role in the physiology of GABAergic neurons (47). Notably, anti-GAD autoantibodies are present not only in type 1 and 2 diabetes mellitus, but also in various neurological diseases including stiff-person syndrome, Miller Fisher syndrome, limbic encephalopathy, cerebellar ataxia, eye movement disorders, and epilepsy, but usually not in MS (48). Collectively, aberrant T cell-mediated autoimmunity against these endogenous self-antigens may disrupt the physiological integrity of the CNS microenvironment and thereby contribute to the pathogenesis of MS. Moreover, these findings underscore the potential utility of an unbiased computational screen in search of etiologically relevant epitopes contributing to the onset and/or aggravation of various autoimmune diseases.

The general principles of Treg-inducing epitopes have remained largely undefined. Treg-inducing epitopes identified from IgG are well-known examples (49, 50), which has been thought to at least partially explain the beneficial immunomodulatory effects of high-dose intravenous immunoglobulin (IVIg) therapy for some autoimmune and autoinflammatory diseases (51). However, the immunomodulatory effect of IgG-derived epitopes is apparently not disease-specific. Identification of disease-specific, etiologically relevant novel Treg-inducing epitopes would provide valuable insights into the pathophysiological mechanisms of various autoimmune diseases, and may also pave the way toward epitope-specific immunomodulatory therapeutics (5, 9, 52). Notably, we found that the majority of the HLA-I predisposing epitopes, although apparently derived from non-self proteome, do not have any positive T cell assay annotation in IEDB. This may be explained as a reflection of tolerance mechanisms leading to clonal anergy or deletion, although overinterpretation should be avoided due to the retrospective nature of this study. In a case study of MS-associated epitopes, we noted that the vast majority of HLA-I predisposing epitopes were derived from HERV-W. HERV-W has also been called MS-associated retrovirus (MSRV) and extensively studied as a potential etiology of MS (53). A recent meta-analysis showed a strong association between detectable HERV-W mRNA and MS (54). Since HERV had been integrated into the human genome in 70 to 30 million years ago, representing almost 8% of the entire human genome, it is not surprising that our adaptive immunity tolerates HERV proteins. Indeed, recent studies highlighted the indispensable role of CD8^+^ Tregs, as well as conventional CD4^+^Foxp3^+^ Tregs, both in the maintenance of self-tolerance and suppression of antiviral immunity (55, 56). Likewise, our analysis of molecular mimicry epitopes revealed human and mycobacteria HSPs as a source of RA-protective HLA-II epitopes, which is consistent with previous observations that HSP-derived epitopes induce CD4^+^ Tregs and provide protection against adjuvant-induced arthritis in mice (20, 38). Therefore, a set of HLA-I non-self epitopes with high DPI but without evidence of T cell recognition, and a set of HLA-II self-epitopes with low DPI, may be considered a good starting point in the exploration of Treg-inducing epitopes in a disease-specific context. Further research is warranted to test the generalizability of this concept.

Molecular mimicry has long been suspected as a general principle of the etiology of various autoimmune diseases. A critical obstacle in the research of molecular mimicry is, in our opinion, that whether any pair of self-and pathogen-derived epitopes have etiological relevance in the same direction, *i.e.*, predisposition to or protection against disease, cannot be argued either based on sequence homology or recognition by the same T cell clones alone. Our framework thus provides one additional layer of criteria for molecular mimicry. It is notable that only four among 43 pairs of self-and pathogen-derived epitopes met the new criteria, which indeed implies that sequence-homologous human and pathogen-derived epitopes could have a distinct impact in terms of disease initiation. This larger-than-expected dissociation between self-epitopes and homologous pathogen-derived epitopes needs to be tested using experimental animal models. In particular, it may be interesting to investigate the putatively disease-predisposing pathogen-derived epitopes homologous to putatively disease-protective self-epitopes. This is because the adaptive immunity in individuals with disease-predisposing HLA alleles may be naïve to the self-epitopes that are likely to be abundantly presented on disease-protective HLA molecules (*i.e.*, potentially tolerance-inducing), and therefore may not be able to tolerate the homologous pathogen-derived epitopes suddenly beginning to be presented upon infection/reactivation of the pathogen(s). Therefore, we would like to propose that etiological roles of pathogen-derived mimicking epitopes should be investigated in a fashion that integrates genetic association, HLA binding, and T cell recognition both *in vitro* and *in vivo*.

The caveats of our concept of differential presentation-based epitope prioritization are as follows. First, the association between disease status and HLA loci does not guarantee an etiological involvement of the encoded HLA molecules, due to the possibility of linkage disequilibrium. Second, affinity to HLA is not the sole parameter explaining pathogenicity (12). For example, the expression level and/or stability of HLA molecules can matter; for example, the low stability of HLA-DQ6 is thought to confer protection against various autoimmune diseases. Moreover, entirely different immunological consequences (*i.e.*, induction of cytopathic vs. regulatory T cells) resulting from alternative TCR docking due to different peptide register have been reported (15). Third, sequence-based affinity prediction may not always reflect the *bona fide* affinity *in vivo*. A good example is post-translationally modified epitopes such as citrullinated epitopes bound to RA-predisposing HLA-DR4 (57). Therefore, further research including experimental validation both *in vitro* and *in vivo* is necessary to comprehensively characterize the HLA-peptidome landscape across autoimmune diseases.

## 4 Materials and Methods

### Computational analysis

All computational analyses were conducted using R ver. 3.5.2 (https://www.r-project.org/) (58). The latest versions of R packages were consistently used. Compiled datasets and essential in-house functions are available as the R package *DPA* on GitHub (https://github.com/masato-ogishi/DPA). Full analytical scripts are available upon request.

### Disease-HLA allele association

Associations between autoimmune diseases and HLA alleles were extracted from a previous PheWAS study (11). This study involved two populations of European ancestry individuals (N=28,839 and 8,431) and tested the association of HLA variation with 1,368 phenotypes. The following autoimmune disease phenotypes were identified: ankylosing spondylitis (AS), celiac disease (CD), dermatomyositis (DM), giant cell arteritis (GCA), Graves’ disease (GD), Juvenile rheumatoid arthritis (JRA), localized lupus and systemic lupus erythematosus (SLE), multiple sclerosis (MS), polymyalgia rheumatica (PMR), polymyositis (PM), primary biliary cirrhosis (PBC), psoriasis (including psoriasis and related disorders, psoriasis vulgaris, and psoriatic arthropathy) (PSO), rheumatoid arthritis (RA), systemic sclerosis (SS), type 1 diabetes (T1D) (including T1D with ketoacidosis and with neurological/ophthalmic/renal manifestations), ulcerative colitis (UC), and Wegener’s granulomatosis (GPA). Only HLA alleles with *P* values of less than 0.01 were included in the analysis. HLA alleles with odds ratios (ORs) of higher and lower than 1 were considered disease-predisposing and disease-protective, respectively. Only autoimmune diseases having both predisposing and protective alleles were included in the subsequent analysis. Disease-HLA associations are summarized in Table S1.

### Epitope sequence datasets

HLA-I-restricted epitope sequences of 8-aa to 14-aa lengths and HLA-II-restricted epitope sequences of 9-aa to 32-aa lengths previously studied in the context of autoimmunity with annotations of functional T cell assay results were collected from Immune Epitope Database (IEDB, as of November 19^th^, 2018) (22). Inclusion/exclusion criteria in terms of T cell assay annotation were provided in Table S2. Epitopes studied in non-human hosts were excluded. Post-translational modifications of epitope sequences were not considered. Epitopes studied in the context of the following autoimmune diseases were identified: AS, CD, GD, GPA, MS, PBC, PSO, RA, SLE, SS, and T1D. Disease-epitope associations and accompanying annotations are summarized in Table S3.

### HLA binding prediction

All four-digit HLA alleles significantly associated with a specific autoimmune disease *X* were used for HLA binding prediction of an epitope *Y* that has been studied in the context of *X*. NetMHCpan 4.0 and NetMHCIIpan 3.2 were utilized for HLA binding prediction with default parameter sets (16, 17). For HLA-DP and DQ alleles, the prediction was conducted for all combinations between the disease-associated target allele and all available counterparts. For example, HLA-DQA1*0102 is associated with predisposition to MS. In this case, binding was predicted against all possible combinations of HLA-DQA1*0102 and HLA-DQB alleles. Then, medians of both predicted affinities and percentile rank values were taken as representative values for the HLA-DQA1*0102 allele.

### Differential presentation analysis

Predicted percentile rank was chosen as a metric for the strength of epitope binding because this metric is not affected by inherent bias of specific HLA molecules towards higher or lower mean predicted affinities and thus allows a direct comparison between different HLA molecules (16). The highest values of sign-inverted log10-transformed percentile ranks, corresponding to the lowest percentile ranks, among predisposing and protective HLA molecules were adopted for differential presentation analysis. Differential presentation index (DPI) was defined as the transformed value of predisposing alleles subtracted by that of protective alleles. DPI is disease-dependent because the sets of predisposing and protective alleles vary between diseases. Epitopes were then categorized in a binary fashion; epitopes with DPI of higher than 0.5 and lower than −0.5 were considered putatively disease-predisposing and putatively disease-protective. Note that epitopes predicted not to bind to any of the disease-associated alleles, with the thresholds being 2% and 10% for HLA-I and HLA-II binding prediction, respectively, were excluded from this binary categorization.

Categorical associations were tested between putative disease association, self/non-self origin, and T cell recognition. Origin was determined by aligning the epitope sequence to the human proteome (UniProt ID: UP000005640). Epitopes with at least one exact match against the human proteome were defined as self-epitopes. T cell recognition (*i.e.*, immunogenicity) was determined from the IEDB annotation. The existence of at least one qualitatively positive functional T cell assay result was considered evidence of immunogenicity. Note that non-functional assays such as binding to peptide-MHC tetramer were not considered for determining immunogenicity. Inclusion/exclusion criteria in terms of T cell assay annotation were provided in Table S2.

Similarity-to-self was measured by aligning the peptide sequence against the entire human proteome (UniProt ID: UP000005640). Pairwise sequence alignment was performed using the *pairwiseAlignment* function implemented in the *Biostrings* package (59). Smith-Waterman local alignment algorithm was employed, with the substitution matrix, gap-opening cost, and gap-extension cost being PAM30, 9, and 1, respectively. These parameters are identical to those utilized in the blastp-short program (see also the BLAST Command Line Applications User Manual: https://www.ncbi.nlm.nih.gov/books/NBK279684/). The highest alignment score among the human protein sequences was used as a metric of similarity-to-self. Likewise, similarity to MBP was defined as the alignment score against human MBP (UniProt ID: P02686) using the same alignment strategy.

## 5 Acknowledgments

The author thanks Dr. Mai Yamakawa and Dr. Wataru Otsu for helpful discussions.

## 7 Author Contributions

M.O. conceived the concept; M.O. performed computational analyses; M.O. wrote the manuscript.

## 9 Data Availability Statement

The datasets and in-house codes necessary to reproduce the work are available as the R package *DPA* on GitHub (https://github.com/masato-ogishi/DPA/). Full analytical scripts are available upon request.

## Supplementary Material

## Supplementary Figures

## Supplementary Tables (Separate files)

**Table S1.** A summary of disease-HLA allele associations.

**Table S2.** Inclusion/exclusion criteria for T cell assay annotations.

**Table S3.** A summary of epitopes previously studied in the context of autoimmune diseases.

**Table S4.** The results of differential presentation analysis in autoimmunity-associated epitopes.

**Table S5.** The results of MS-specific differential presentation analysis in self-epitopes previously studied in various contexts.

**Table S6.** The results of differential presentation analysis in self-epitopes and corresponding pathogen-derived epitopes with evidence of molecular mimicry.

**Figure S1.**
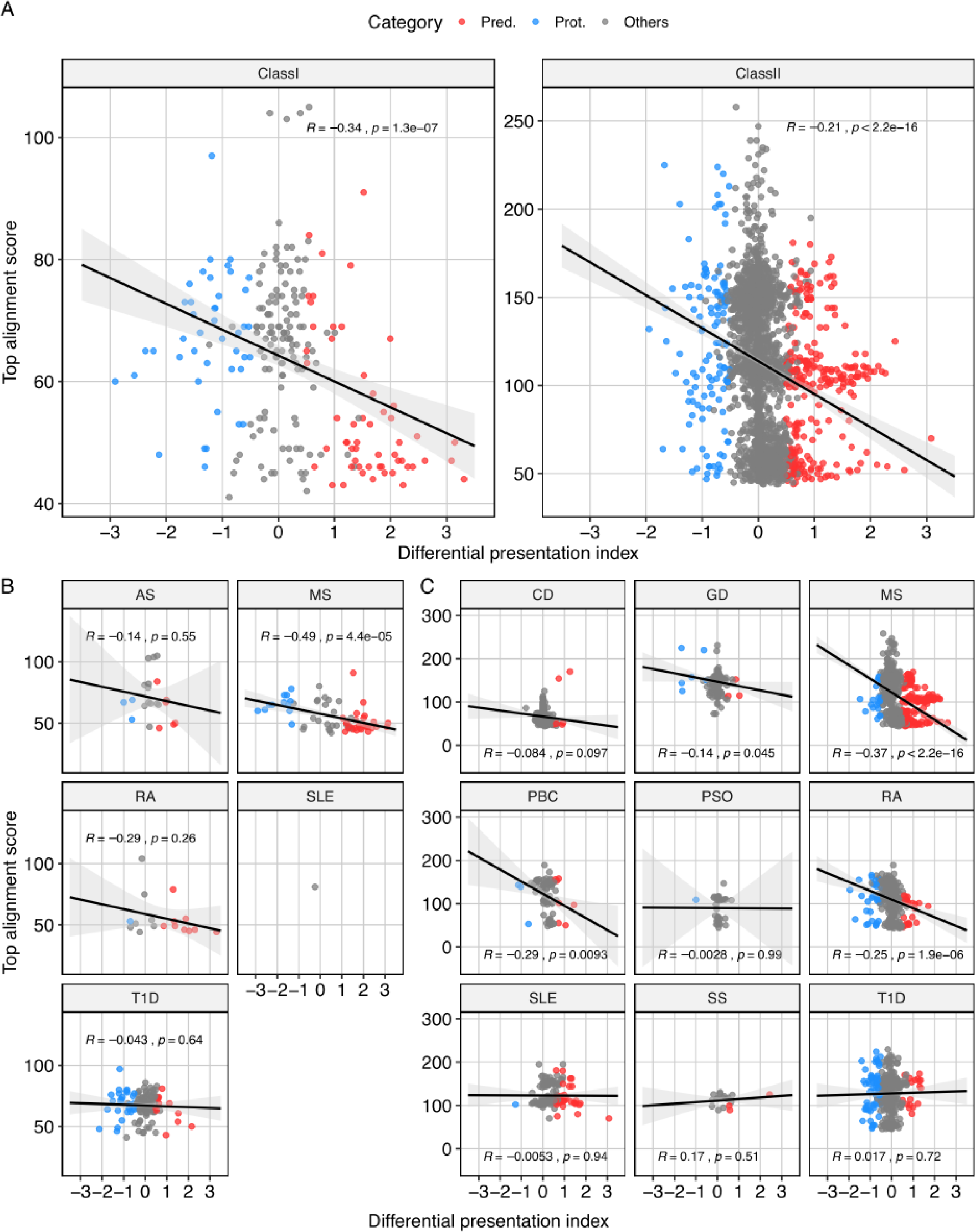
Correlations between DPI and similarity-to-self. (A) DPI-selfness two-dimensional plots of autoimmunity-associated HLA-I and HLA-II epitopes. Autoimmunity-associated epitopes were analyzed, and disease-specific DPI scores were calculated. The highest alignment score against the human proteome (UniProt ID: UP000005640) was also computed for each of the epitopes. (B and C) Two-dimensional plots of (B) HLA-I and (C) HLA-II epitopes facetted by the associated diseases. *R* indicates Pearson’s correlational coefficient. For disease abbreviations, see Figure 2.

**Figure S2.**
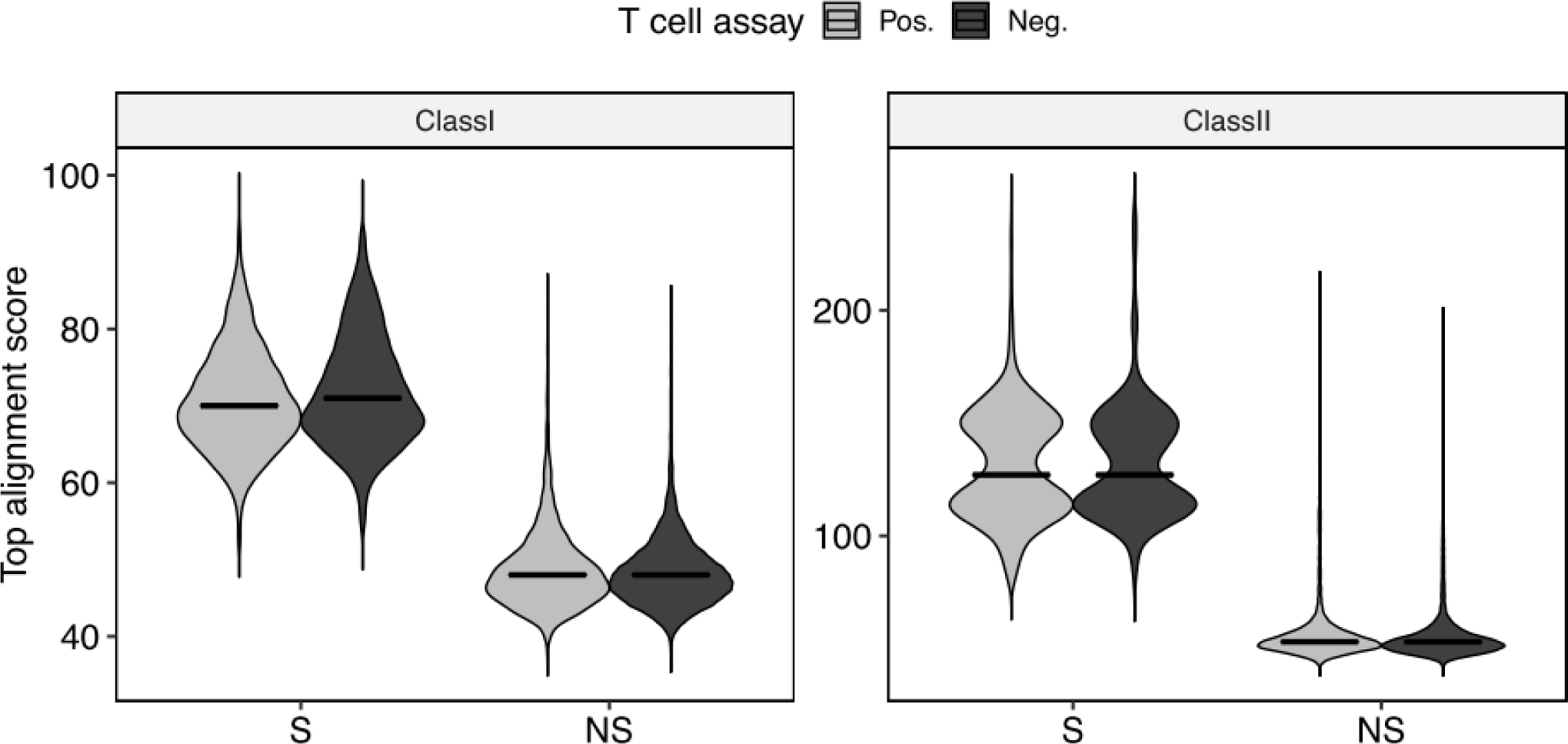
Minimal differences of similarity-to-self among immunogenic and non-immunogenic epitopes. Epitopes studied in various contexts were compiled previously. Epitopes of self (S) and non-self (NS) origins were identified based on the presence or absence of at least one exact sequence match in the human proteome. Immunogenicity was determined based on the presence or absence of at least one positive T cell assay annotation.

